# Cleaning genotype data from Diversity Outbred mice

**DOI:** 10.1101/518308

**Authors:** Karl W. Broman, Daniel M. Gatti, Karen L. Svenson, Śaunak Sen, Gary A. Churchill

## Abstract

Data cleaning is an important first step in most statistical analyses, including efforts to map the genetic loci that contribute to variation in quantitative traits. Here we illustrate approaches to quality control and cleaning of array-based genotyping data for multiparent populations (experimental crosses derived from more than two founder strains), using MegaMUGA array data from a set of 291 from Diversity Outbred (DO) mice. Our approach employs data visualizations that can reveal problems at the level of individual mice or with individual SNP markers. We find that the proportion of missing genotypes for each mouse is an effective indicator of sample quality. We use microarray probe intensities for SNPs on the X and Y chromosomes to confirm the sex of each mouse, and we use the proportion of matching SNP genotypes between pairs of mice to detect sample duplicates. We use a hidden Markov model (HMM) reconstruction of the founder haplotype mosaic across each mouse genome to estimate the number of crossovers and to identify potential genotyping errors. To evaluate marker quality, we find that missing data and genotyping error rates are the most effective diagnostics. We also examine the SNP genotype frequencies with markers grouped according to their minor allele frequency in the founder strains. For markers with high apparent error rates, a scatterplot of the allele-specific probe intensities can reveal the underlying cause of incorrect genotype calls. The decision to include or exclude low-quality samples can have a significant impact on the mapping results for a given study. We find that the impact of low-quality markers on a given study is often minimal, but reporting problematic markers can improve the utility of the genotyping array across many studies.

## Introduction

Data cleaning is a critical first step in analyses to map quantitative trait loci (QTL). Genotyping errors and especially the inclusion of poor quality or erroneously labeled samples can reduce the power to detect QTL. Despite its importance, relatively little has been written about the data cleaning process. Lincoln and Lander (1992) discussed the detection of genotyping errors in genetic map construction. Broman and Sen (2009, Ch. 3) discussed data cleaning more broadly, in biparental crosses. Morgan (2015), in presenting the R package argyle for quality control for SNP genotyping arrays, emphasized consideration of the amount of missing genotypes, frequency of heterozygotes, and the distribution of array intensities. Here we address data cleaning with an emphasis on multiparent populations and array-based genotyping.

Multiparent populations are experimental crosses derived from more than two founder strains. They have become a popular tool for complex trait genetics in experimental organisms. Examples include heterogeneous stock (Mott *et al*. 2000; Mott and Flint 2002), MAGIC lines (Cavanagh *et al*. 2008; Kover *et al*. 2009), the Collaborative Cross (Churchill *et al*. 2004), and Diversity Outbred mice (Churchill *et al*. 2012; Svenson *et al*. 2012). Genotype data cleaning is more difficult in multiparent populations as individual SNP markers are generally limited to two alleles and thus multiple marker genotypes are required to uniquely identify the founder strain origins at any locus.

We illustrate our process for cleaning genotype data using MegaMUGA SNP array data (Morgan *et al*. 2016) on 291 Diversity Outbred (DO) mice. The SNP probe selection for this and other MUGA platforms has been optimized to distinguish among the founder haplotypes of the DO. A key principle that guides our approach to data cleaning is to think about what might have gone wrong, and how it might be revealed in the data. We visualize the data in many ways, and when we see something unexpected, we try to determine the underlying cause.

## Material and Methods

### Mice and genotypes

We consider genotype data for 291 Diversity Outbred (DO) mice, including 99 mice from generation 8 and 192 mice from generation 11. These are a subset of the mice considered in Gatti *et al*. (2017).

The mice were genotyped using the MegaMUGA SNP array (Morgan *et al*. 2016), which includes 77,808 markers. The genotyping was performed at Neogen (Lincoln, NE). Genotype calls using pairs of nucleotides A, C, G, and T were converted to genotypes AA, AB, BB, with A denoting the allele that was most frequent among the eight founder strains (assigned arbitrarily when the two alleles were equally frequent).

### Statistical methods

Analyses were conducted in R (R Core Team 2018) and with R/qtl2 (Broman *et al*. 2019). Most of the statistical analyses involves visualization of summary statistics.

To reconstruct the genomes of the DO mice, we use a hidden Markov model (see Broman and Sen 2009, App. D). The transition probabilities for DO mice were taken from Broman (2012b), which uses the results of Broman (2012a). We used the genetic map from Cox *et al*. (2009), assumed a genotyping error rate of 0.2%, and used the Carter-Falconer map function (Carter and Falconer 1951). The HMM provides a probability for each possible 36-state diplotype at each marker for each mouse. The term diplotype refers to a pair of founder haplotypes. The 36 diplotypes consist of 8 homozygotes and 28 heterozygotes.

To estimate the number of crossovers in each individual, we inferred the 36-state diplotype at each locus as the state with maximum marginal probability, provided that it was > 0.5. In cases where all diplotypes had probability < 0.5, the diplotype state was treated as missing. We then calculated the minimum number of crossovers consistent with these inferred diplotypes.

To identify potential genotyping errors, we calculated the genotyping error LOD scores of Lincoln and Lander (1992), by first converting the 36-state diplotype probabilities to 3-state SNP genotype probabilities, using the SNP genotypes in the eight founder strains. We then used the observed SNP genotype to calculate the error LOD score as in equation 1b of Lincoln and Lander (1992).

To obtain predicted SNP genotypes for each DO mouse, we collapsed the predicted 36-state diplotypes to 3-state SNP genotypes using the SNP genotypes in the founders. The predicted genotypes can differ from the observed genotypes due to the smoothing effects of the HMM haplotype reconstruction.

### Data and software availability

The raw genotype data are available at FigShare (https://doi.org/10.6084/m9.figshare.7359542.v1). They are also available in the R/qtl2 input format at https://github.com/rqtl/qtl2data.

The SNP genotypes for the founder strains are at FigShare, https://doi.org/10.6084/m9.figshare.5404750.v2. We used annotations for the MegaMUGA genotyping array from https://github.com/kbroman/MUGAarrays.

R/qtl2 is available at https://kbroman.org/qtl2 and at GitHub, https://github.com/rqtl/qtl2. The custom R scripts used for our analyses and to create the figures are at GitHub (https://github.com/kbroman/Paper_MPPdiag).

## Results

We consider data from a population of 291 Diversity Outbred (DO) mice (Gatti *et al*. 2017), including 99 mice from generation 8 and 192 mice from generation 11. There are 150 females and 141 males. The mice were genotyped with the MegaMUGA array, which includes 77,808 markers, but we focus on the 69,339 markers that are polymorphic among the eight founder strains.

### Sample diagnostics

#### Missing data

Our first step in genotype data diagnostics is to look at the proportion of missing genotypes in each mouse, as this is a key indicator of sample quality. Samples with a high proportion of missing genotypes are likely low quality.

The percent missing genotypes by mouse are displayed in Figure 1. The bulk of mice are missing very little data, but there are 24 mice with >2% missing genotypes, including nine mice with ≥20% missing. The mice with high rates of missing genotypes will show up as outliers for many of the diagnostics below. The problem samples are clustered within a common batch, which may indicate a problem with sample processing.

**Figure 1.**
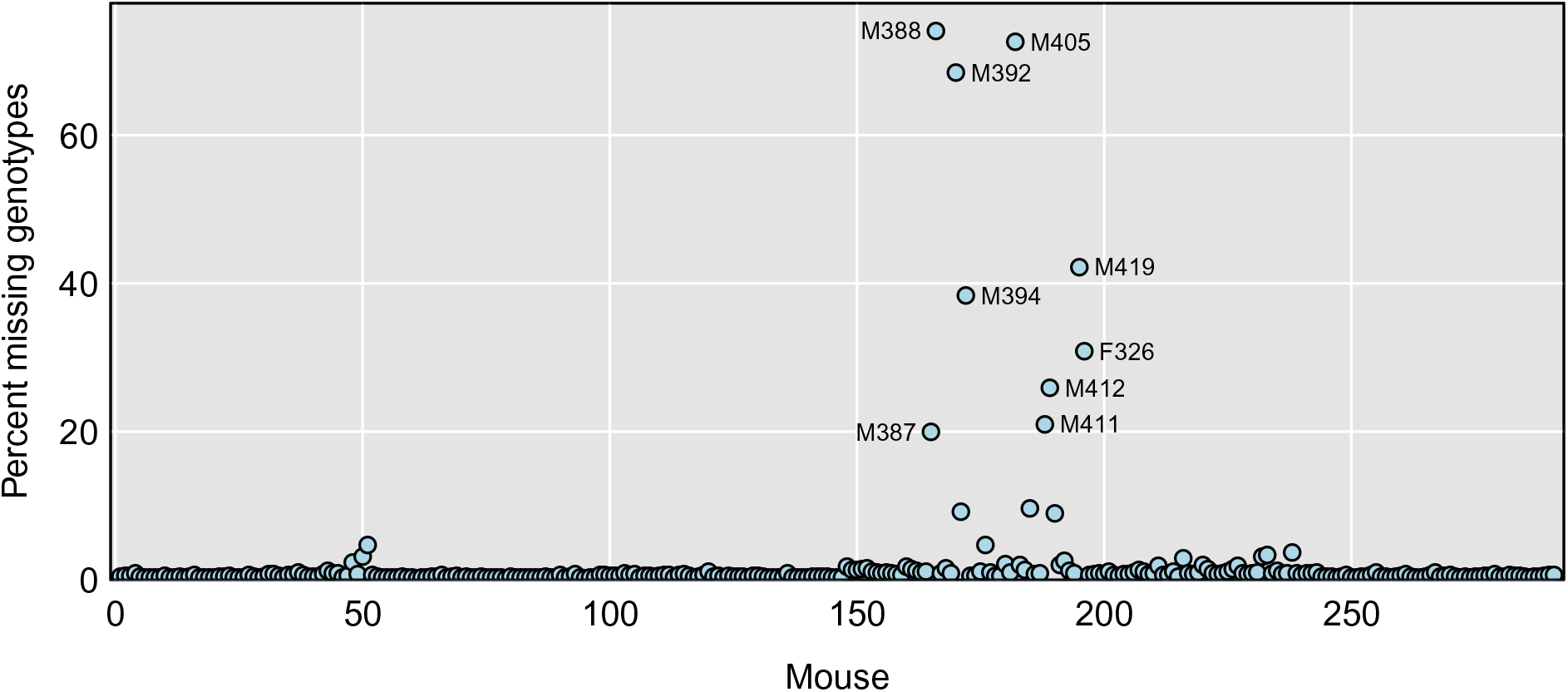
Percent missing genotypes by mouse. The nine mice with ≥20% missing genotypes are labeled with their sample identifiers.

#### Verify sex

To verify the sexes of the mice, we could look at the proportion of heterozygous genotype calls on the X chromosome. However, with array-based genotypes, it is most informative to look at the probe intensities for SNPs on the X and Y chromosomes.

In Figure 2, we display a scatterplot of the average intensity for the 30 Y chromosome markers vs the average intensity for 2,058 X chromosome markers. Each point is a mouse, with the males in purple and the females in green. We omitted eighteen markers on the X chromosome that did not show a clear sex difference in allele intensities.

**Figure 2.**
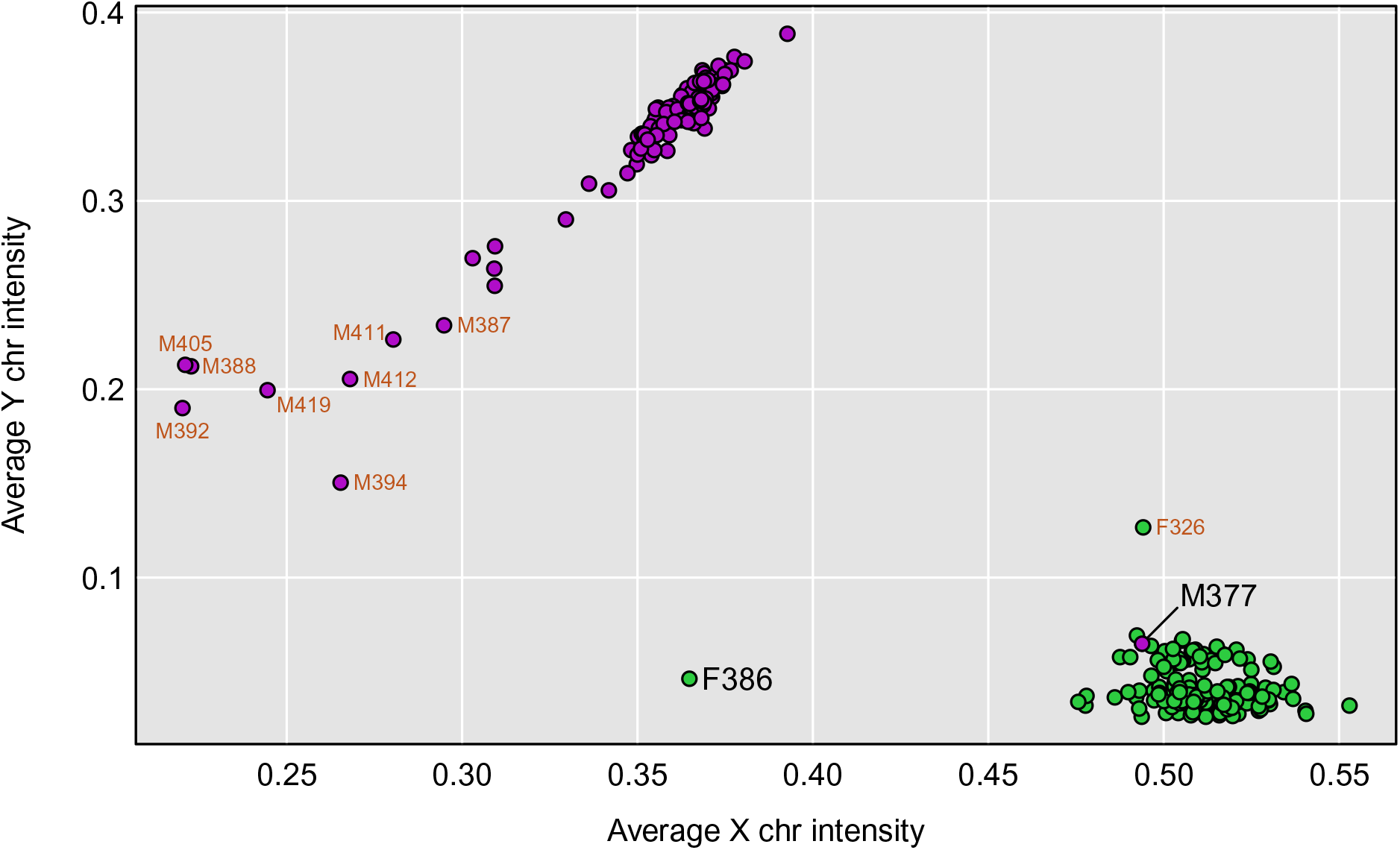
Average SNP microarray intensity for markers on the Y chromosome versus that for markers on the X chromosome, for each mouse. Mice that were nominally male are in purple, while females are in green. Samples with ≥ 20% missing genotypes are labeled in orange. Two other samples of interest are labeled in black: F386 which appears to be XO, and M377 which was nominally male but appears to be XX.

There is a distinct cluster of male mice in the upper-left (low X chromosome intensity and high Y chromosome intensity) and a cluster of female mice in the lower-right (high X chromosome intensity and low Y chromosome intensity). However, M377 was labeled male but appears within the female cluster in the lower right, and so is likely female. Also, F386 is labeled female but appears in the lower-left, with reduced X chromosome intensity. This is likely an XO female. Other outliers in Figure 2 are mice with high rates of missing genotypes, including the nine mice with ≥ 20% missing genotypes, which are labeled in orange.

For sex inference, the SNP probe intensities give better separation of the two sexes than heterozygosity on the X chromosome (Figure S1), and they enable us to distinguish between males and XO females. We could also, potentially, identify XXY males, who would have high average intensities on both sex chromosomes.

#### Sample duplicates

We look for potential sample duplicates by calculating the proportion of matching SNP genotypes for each pair of samples. The nine samples with ≥ 20% missing genotypes look quite different from others, and also show some chance similarities (Supplementary Figure S2A). If we omit those samples (Supplementary Figure S2B), we find that the bulk of pairs share around 50% of genotypes (shifted slightly below 50%; the median is 46.6%). The probability of unrelated individuals sharing the same genotypes at a marker is determined by the SNP minor allele frequency. The average proportion of sharing is a property of both the population structure and the array probe selection strategy. A small proportion of pairs (179 pairs, or 0.4%) share a bit more, at around 67.1%. These likely represent siblings.

Two pairs have almost perfectly matching SNP genotypes. Mice M283 and M292 have matching genotypes at all except one of the 69,025 markers at which they were both genotyped. Mice M377 and F409 have matching genotypes at all except 36 of the 68,291 markers at which they were both genotyped. Note that the second pair includes M377 which was seen in Figure 2 to have mislabeled sex.

These two pairs are clear duplicates. We are looking for a separation between the normal sharing between mice and the duplicates, and (having excluded samples with ≥ 20% missing genotypes), these are the only pairs with >76% matching genotypes. These unintended sample duplicates provide an estimate of the genotyping error rate, which looks to be well under 1/10,000.

We will omit one mouse from each pair, for the purpose of illustrating our genotype quality control analyses. For the M377/F409 pair, it seems clear that the sample corresponds to F409, since it is a female. But for M283/M292, we can not tell which is the correct label. For later QTL analyses we would likely wish to omit both samples, but for this illustration we will just omit the second one, M292.

If the data included genome-scale phenotypes with strong genetic signals, such as gene expression data, we would at this point look further for possible sample mix-ups (Westra *et al*. 2011; Broman *et al*. 2015), but we will not do so here.

#### SNP probe intensities

The distribution of probe intensities on the genotyping array can be a useful indicator of problem samples. In Figure 3A, we display density estimates of the array intensities for each of the 289 samples, after a log_10_(*x* + 1) transformation. We highlight the arrays corresponding to samples with appreciable missing genotypes. The nine samples with ≥ 20% missing data are highlighted in orange; their array intensities are shifted to the left and have a long right tail. There are three samples with ∼9% missing data (in pink); they show a much broader distribution of array intensities. There are eight samples with 2.5–5% missing genotypes (in blue); many of these have a spike of intensities near 0.

**Figure 3.**
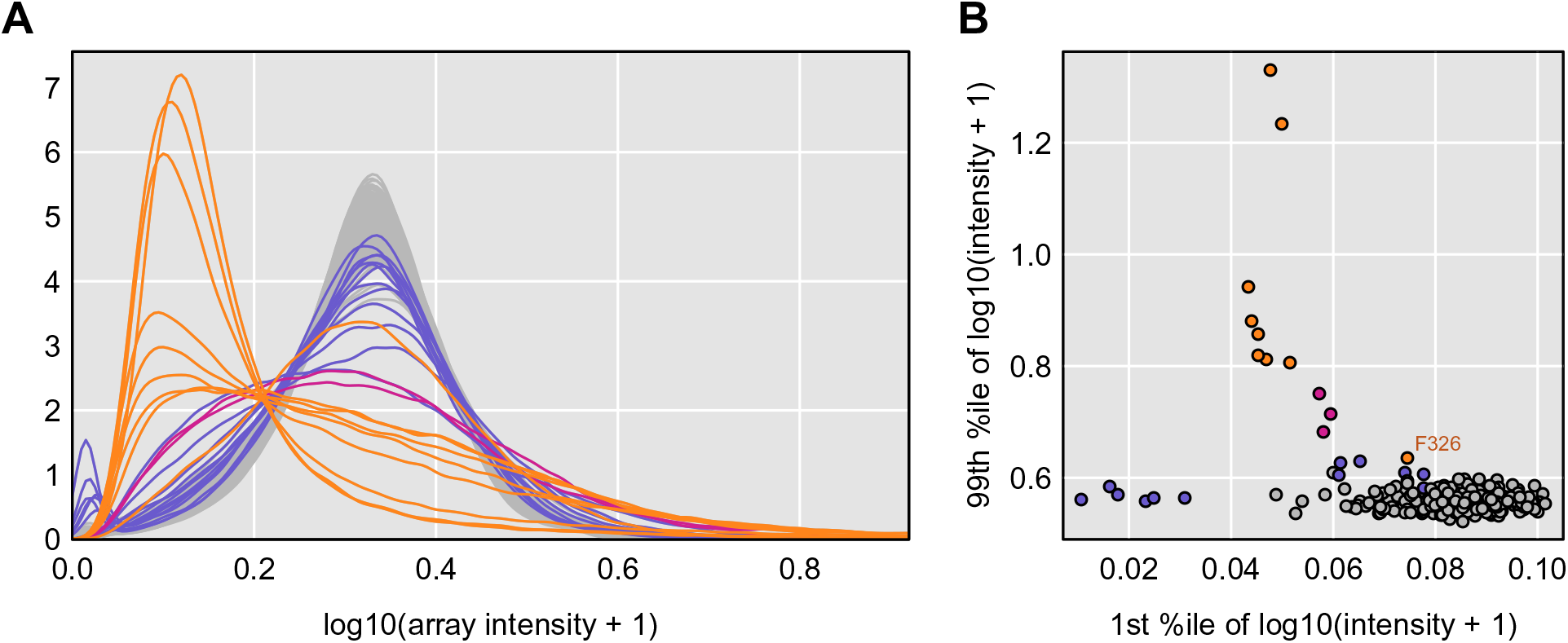
Distribution of array intensities after a log_10_(*x* + 1) transformation. **A**: Kernel density estimates of the array intensity distribution. Samples with >9% missing genotype data are in pink; samples with 2.5 – 5% missing genotype data are in blue; the remaining samples are in gray. **B**: Scatterplot of the 1st percentile versus the 99th percentile.

In Figure 3B, we display a scatterplot of the 1st and 99th percentiles of the log array intensities, in an attempt to summarize the pattern seen in the densities in Figure 3A. Most of the samples with high rates of missing genotypes are outliers. We label one sample (F326) which has ≥20% missing genotypes but has an array intensity distribution that is not as extreme as the other samples with ≥20% missing genotypes

#### SNP genotype frequencies

The SNP genotype frequency distribution across all markers within an animal can be a useful diagnostic. For example, sample contamination can lead to excess heterozygotes. In DO mice, we split the SNP markers into four groups, based on the minor allele frequency (MAF) in the eight founder strains. To do so, we consider only the 68,357 with complete founder genotypes. (For 982 of the polymorphic markers, at least one of the founders has missing genotype.)

To display the genotype frequencies, we use a ternary plot, which makes use of the fact that for any point within an equilateral triangle, the sum of the distances to the three sides is constant. And so the trinomial SNP genotype frequencies for an individual may be represented by a point in the triangle, with the distance to the lower edge being the frequency of heterozygotes, and the distances to the left and right edges being the frequencies of the two homozygotes.

As seen in Figure 4, the DO mice form tight clusters with very similar SNP genotype frequencies, except for a few samples that show high heterozygosity, which are among the samples with high rates of missing genotype data. Another potential outlier is F313, which shows somewhat reduced heterozygosity in SNPs with MAF = 3/8 or 1/2.

**Figure 4.**
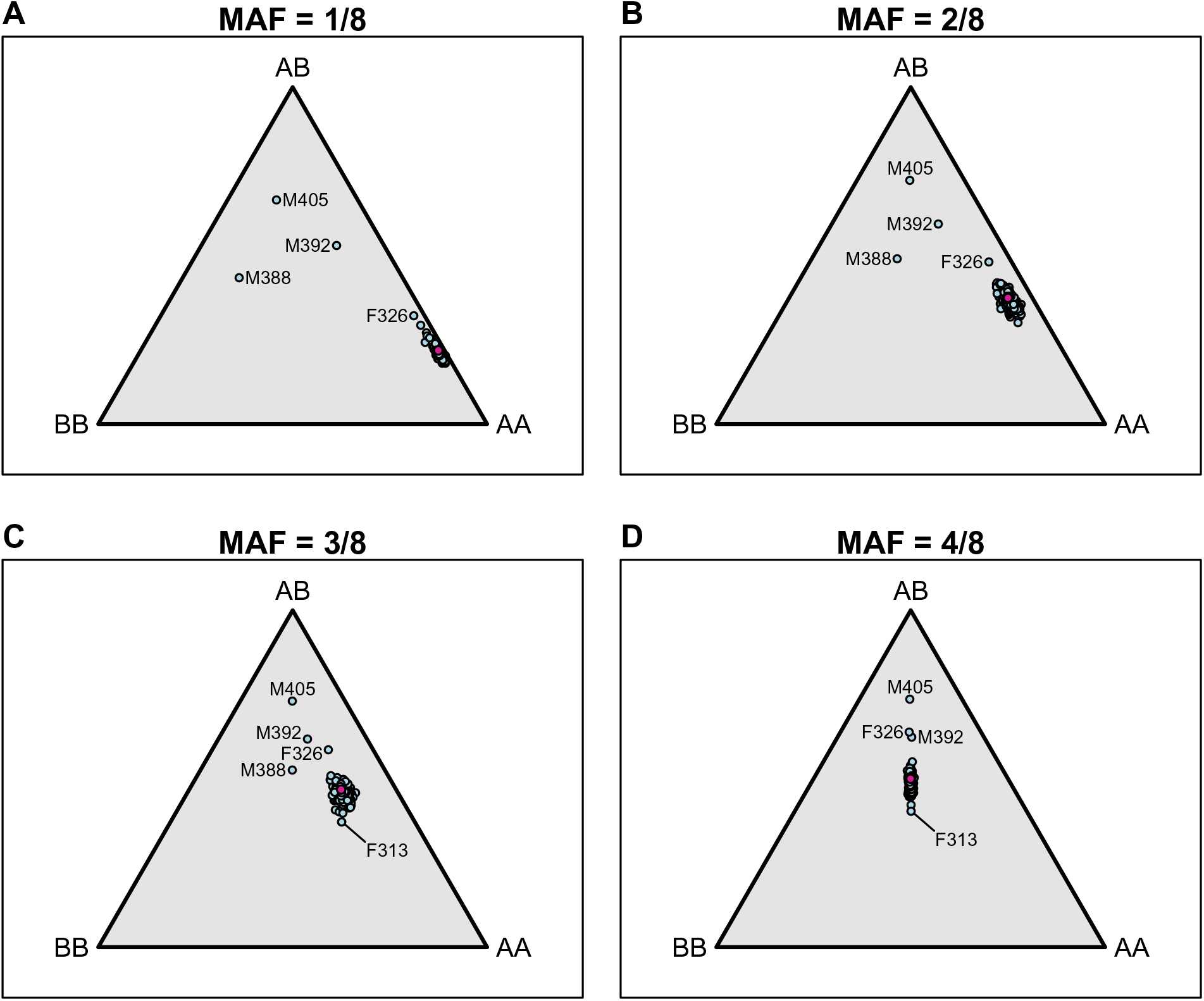
SNP genotype frequencies by mouse, for SNPs split by their minor allele frequency (MAF) in the eight founder strains. Trinomial probabilities are represented by points in an equilateral triangle using the distances to the three sides. Pink points indicate the expected distributions.

#### Counts of crossovers

Another important diagnostic is to estimate the number of crossovers, genome-wide, in each mouse. Problem samples may exhibit excessive crossovers. In some cases, a sample may exhibit fewer crossovers than expected.

To estimate crossovers, we first reconstruct the diplotypes across the genome for each of the DO mice. We use a hidden Markov model (HMM) to calculate the probability of each of the 36 possible diplotypes (8 homozygotes plus 28 heterozygotes) at each genotyped SNP, given the multipoint SNP genotype data, with allowance for genotyping errors (Supplementary Figure S3). Figure S3A shows the inferred diplotypes across the genome for a single DO mouse, and Figure S3B shows the detailed diplotype probabilities for that mouse, along one chromosome.

There are a number of different methods for estimating the number of crossovers in a DO genome. We are using the simplest: at each genotyped SNP, pick the most probable diplotype (provided that it has probability at least 50%) and then calculate the minimal number of crossovers that are consistent with that set of predicted diplotypes.

The estimated numbers of crossovers for each DO mouse is shown in Figure 5, with points colored according to their generation. The mice in generation 8 have an average of 304 crossovers, while those in generation 11 have an average of 357 crossovers.

**Figure 5.**
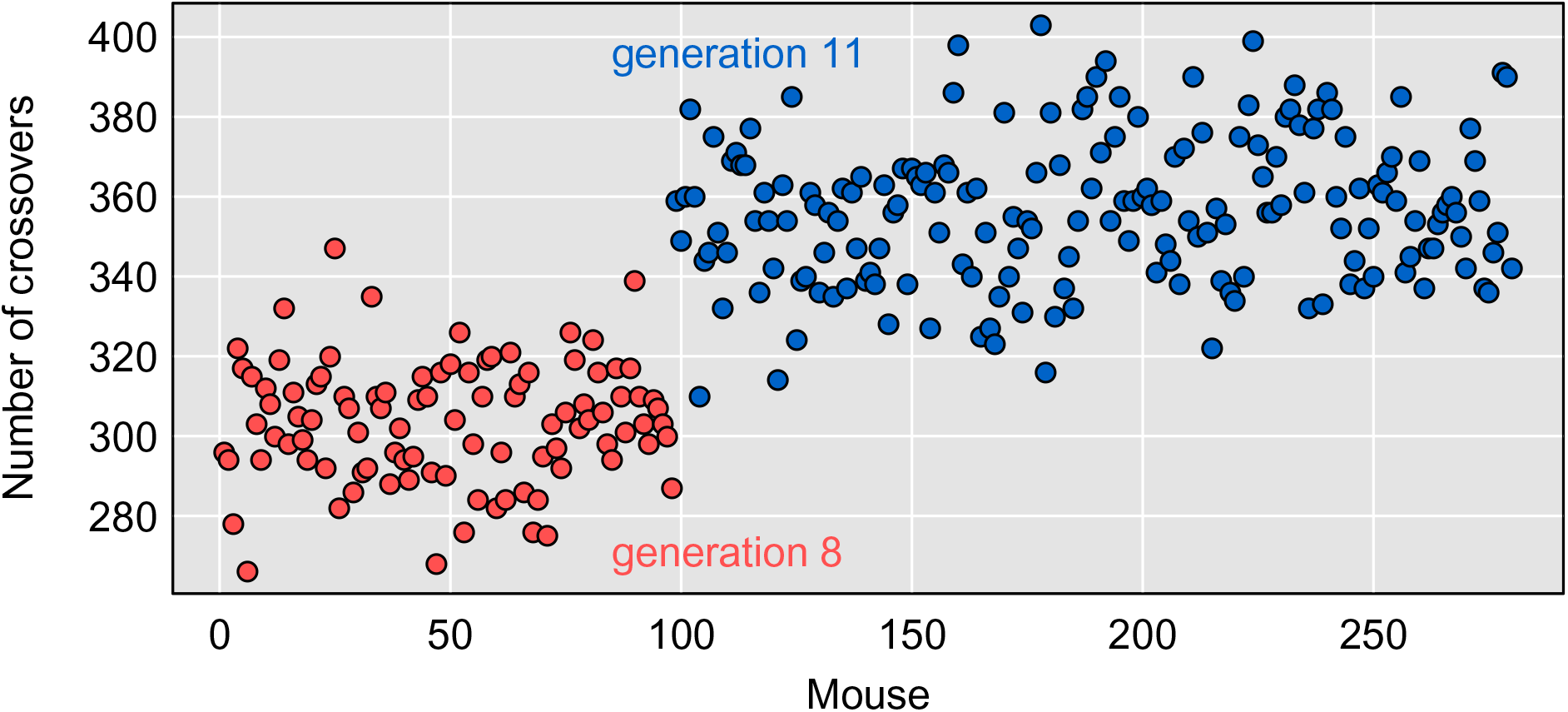
Estimated number of crossovers in each mouse. Colors of the points indicate the two groups of DO mice (generations 8 and 11). Mice with ≥ 20% missing genotypes are excluded.

We have excluded the nine mice with ≥ 20% missing genotypes. They all show > 500 crossovers, and the mice with > 50% missing genotypes show > 2000 crossovers. This is further evidence that these samples should be omitted.

There are no other apparent outliers, in the numbers of crossovers. But note the importance of taking account of generation number.

#### Genotyping error rates

The diplotype reconstructions offer the opportunity to identify likely SNP genotyping errors. To identify likely errors, we calculate the genotyping error LOD scores described by Lincoln and Lander (1992). For each SNP marker in each DO mouse, we use the founder strains’ SNP genotypes to collapse our 36-state diplotype probabilities to three-state SNP genotype probabilities. We then compare the predicted SNP genotypes based on the haplotype reconstruction with the observed SNP genotypes, and we calculate a LOD score statistic that measures the evidence for an individual SNP genotype being in error.

We generally focus on error LOD scores >2, which is a reasonably conservative threshold on potential errors. The estimated error rate in each mouse, with this criterion, is shown in Figure 6. The nine mice with ≥ 20% missing genotypes all have estimated genotyping error rates > 1%. Three mice have error rates in the range 0.5–1.0%, and these all showed ∼9% missing genotypes. The next-highest estimated error rate is 0.4% for mouse M398, which had shown about 5% missing genotypes. The error rates for the other mice are extremely small. The median rate is just 7.8 in 10,000.

**Figure 6.**
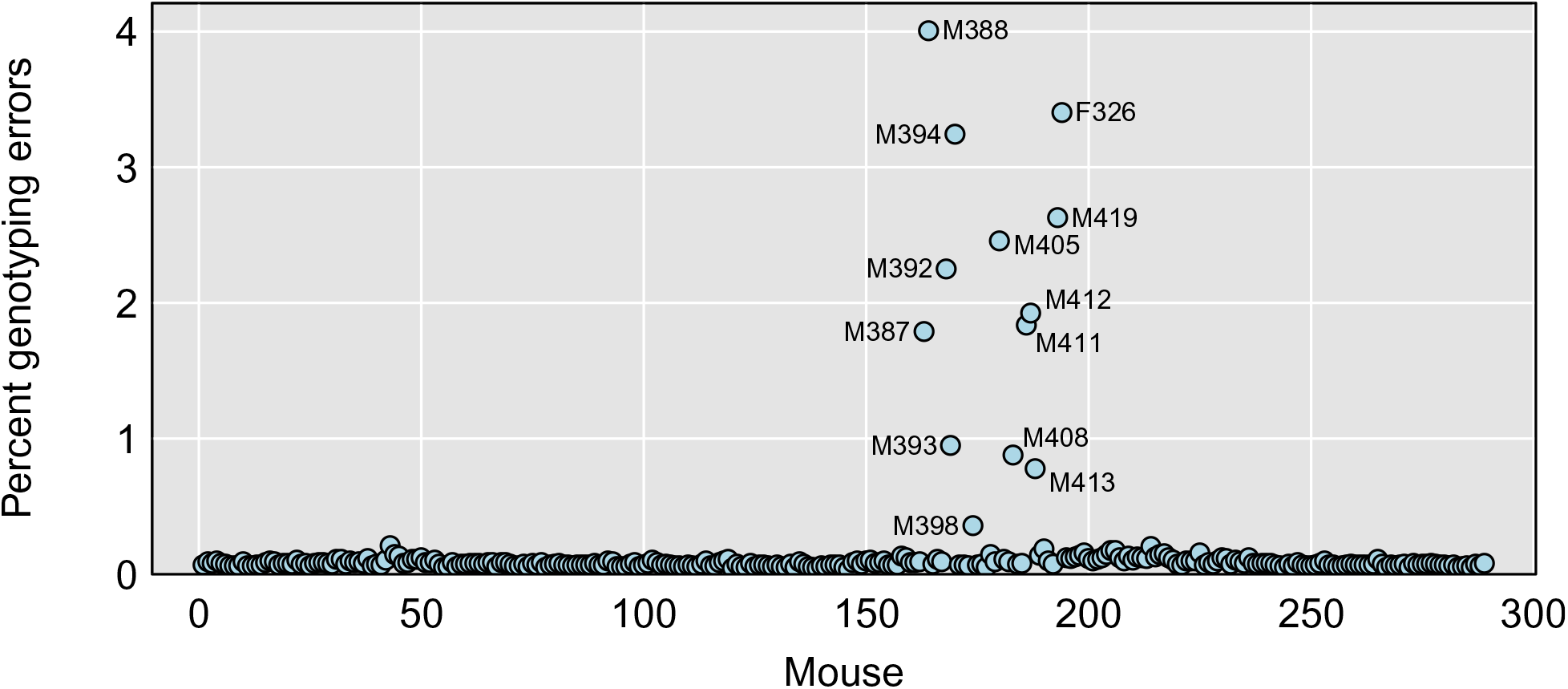
Estimated percent genotyping errors for each mouse. The rates are very small; the median is just 7.8 in 10,000.

In summary, the nine mice with ≥ 20% missing genotypes also showed excessive crossovers and high rates of apparent genotyping errors. We will omit these from further analyses. Three mice with 5–10% missing genotypes showed slightly elevated genotyping error rates but no excess of crossovers. We chose to not omit them.

The factors that impact the decision to omit samples will vary from study to study and may also depend on the aims of the analysis. Samples that are clearly outliers will likely have a negative impact on mapping power and precision. It is not uncommon for some samples to fall into a gray area where the quality is less than ideal but, based on the number of predicted crossovers, the diplotype reconstruction appears to be reliable. The impact on power of omitting these samples may be greater if the sample size is already small.

### Marker diagnostics

We now turn to the markers, to identify poorly behaved ones. As with the samples, we can look at the percent missing data, estimated rates of genotyping errors, and the genotype frequencies. Ultimately, we want to look at scatterplots of the allele-specific probe intensities for the SNPs, which are most informative of problems, but with 77,808 markers, we cannot inspect all of them, and so we use the other measures to help narrow our search. Throughout these analyses, we will focus on the 280 mice with < 20% missing genotypes.

#### Missing data and genotyping errors

We start by studying the amount of missing data at each marker, as well as estimated genotyping error rates (based on genotyping error LOD scores > 2). A scatterplot of estimated error rate vs. percent missing data is shown in Figure 7.

**Figure 7.**
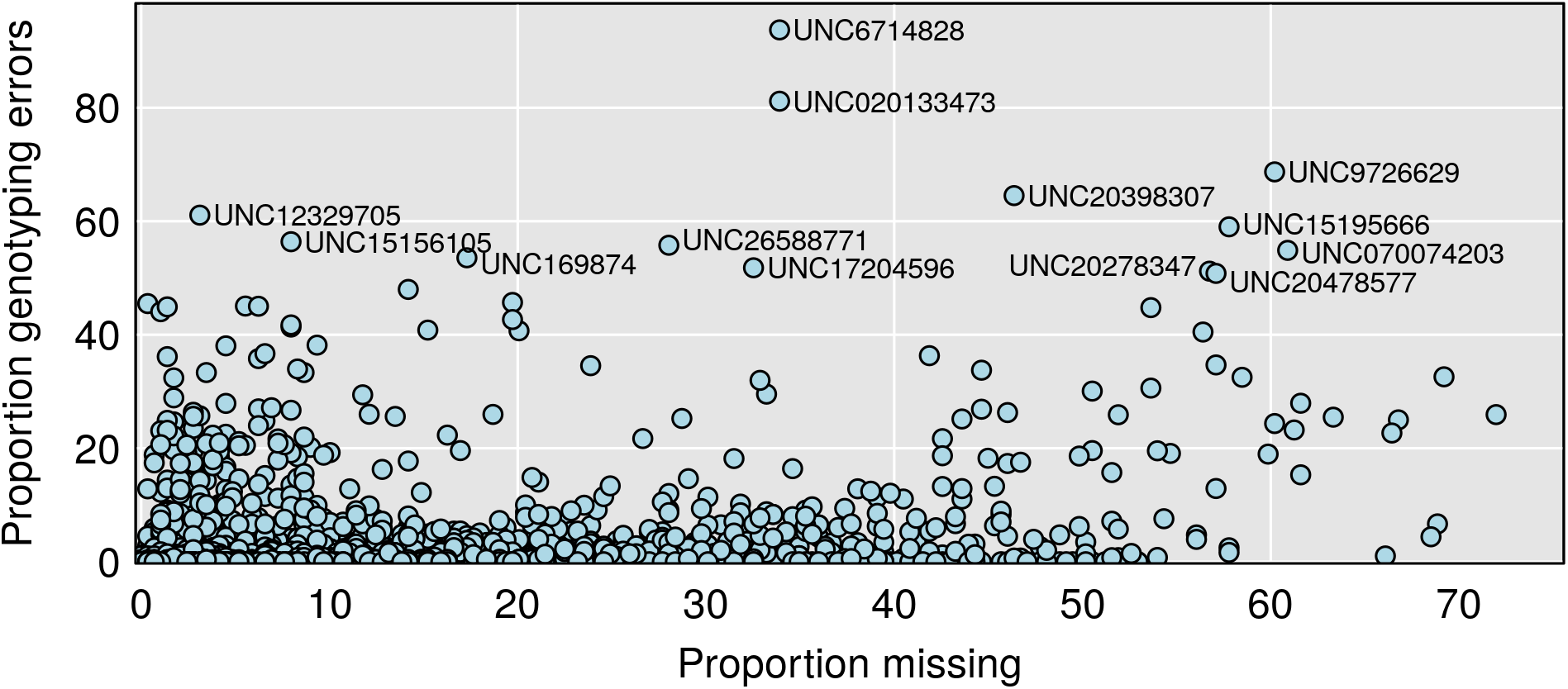
Estimated percent genotyping errors vs. percent missing genotypes by marker. Errors defined by genotyping error LOD score > 2. The vast majority of markers showed no apparent errors.

The vast majority of markers have virtually no missing data: Of the 69,339 informative markers, 41,931 have no missing data, and 64,003 are missing < 2%. However, as seen in Figure 7, there are a number of markers with appreciable missing data: 1,216 are missing > 10%.

Similarly, the vast majority of markers have virtually no genotypes with error LOD score > 2, including 66,623 markers with no apparent errors. But 624 markers have estimated error rates > 2%, including 190 markers with estimated error rates > 10%. The thirteen markers with error rates > 50% are highlighted in Figure 7.

While it may not be apparent in Figure 7, there is a reasonably strong relationship between missing data and error rate: Markers with larger amounts of missing data tend to have higher genotyping error rates. For example, the mean error rate for markers with < 2% missing data is 3 per 10,000, while the mean error rate for markers with ≥ 20% missing data is 5%, about 190 times higher.

#### SNP genotype frequencies

The genotype frequencies at the markers are displayed in Figure 8, with markers split according to their minor allele frequency (MAF) among the 8 founder strains. The majority of markers conform reasonably well to our expectation. The most striking departure is that there are 22 markers with MAF=1/8 in the founders but where the minor allele in the DO mice is > 40%. The majority of these markers (19/22) are from a region on chromosome 2, 70 – 105 Mbp, where the WSB allele exhibits meiotic drive (Didion *et al*. 2016).

**Figure 8.**
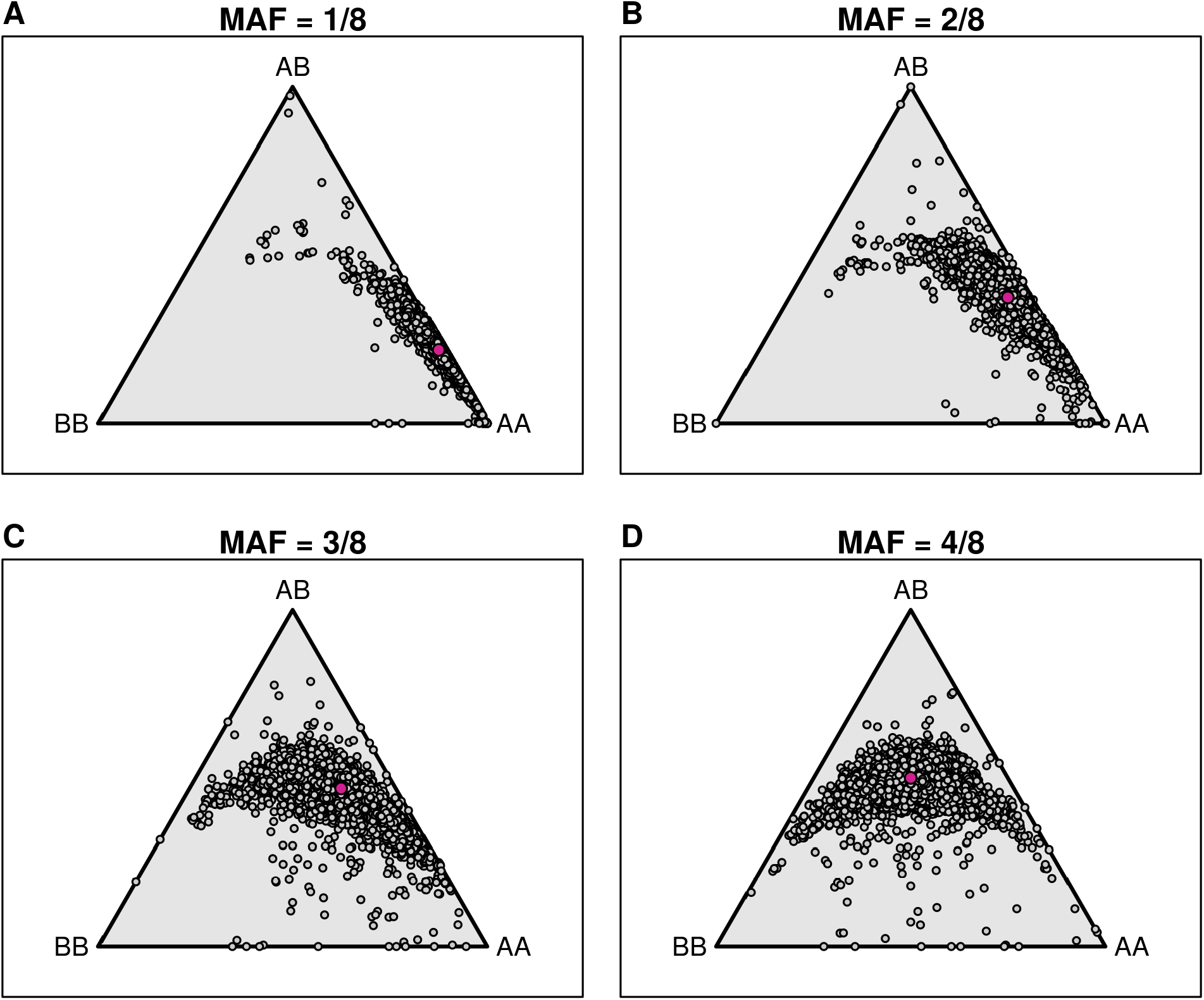
SNP genotype frequencies by marker, with SNPs split by their minor allele frequency (MAF) in the eight founder strains. Trinomial probabilities are represented by points in an equilat­eral triangle using the distances to the three sides. Pink points indicate the expected distributions.

There are also a number of SNPs with MAF = 3/8 or 4/8 in the founder strains that have reduced frequency of heterozygotes. For example, there are 46 SNPs with MAF = 4/8 in the founders but heterozygosity < 0.25. Most of these SNPs (37) have ≥ 10% missing data or ≥ 10% genotyping errors, or both. Of the remaining nine markers, all but one are on chromosome 2, in the region with high WSB allele frequency.

#### SNP allele intensities

The most important diagnostic for SNP quality is a scatterplot of the allele-specific probe intensities. This is particularly informative when colored by both the observed genotype and by the predicted genotype giving the multipoint SNP information. The SNP allele intensities for a set of four SNPs are displayed in Figure 9. Each point is a single DO mouse; in the left panels, the points are colored by the observed genotype, with yellow and blue corresponding to the two homozygotes, green the heterozygote, and gray being missing. In the right panels, the points are colored by the predicted genotypes given the multipoint SNP information. Figures 9A and 9B correspond to a well-behaved SNP. The three genotype groups form tight, well-separated clusters, and the observed and predicted genotypes match. (Additional examples of well-behaved markers are shown in Supplementary Figure S4.)

**Figure 9.**
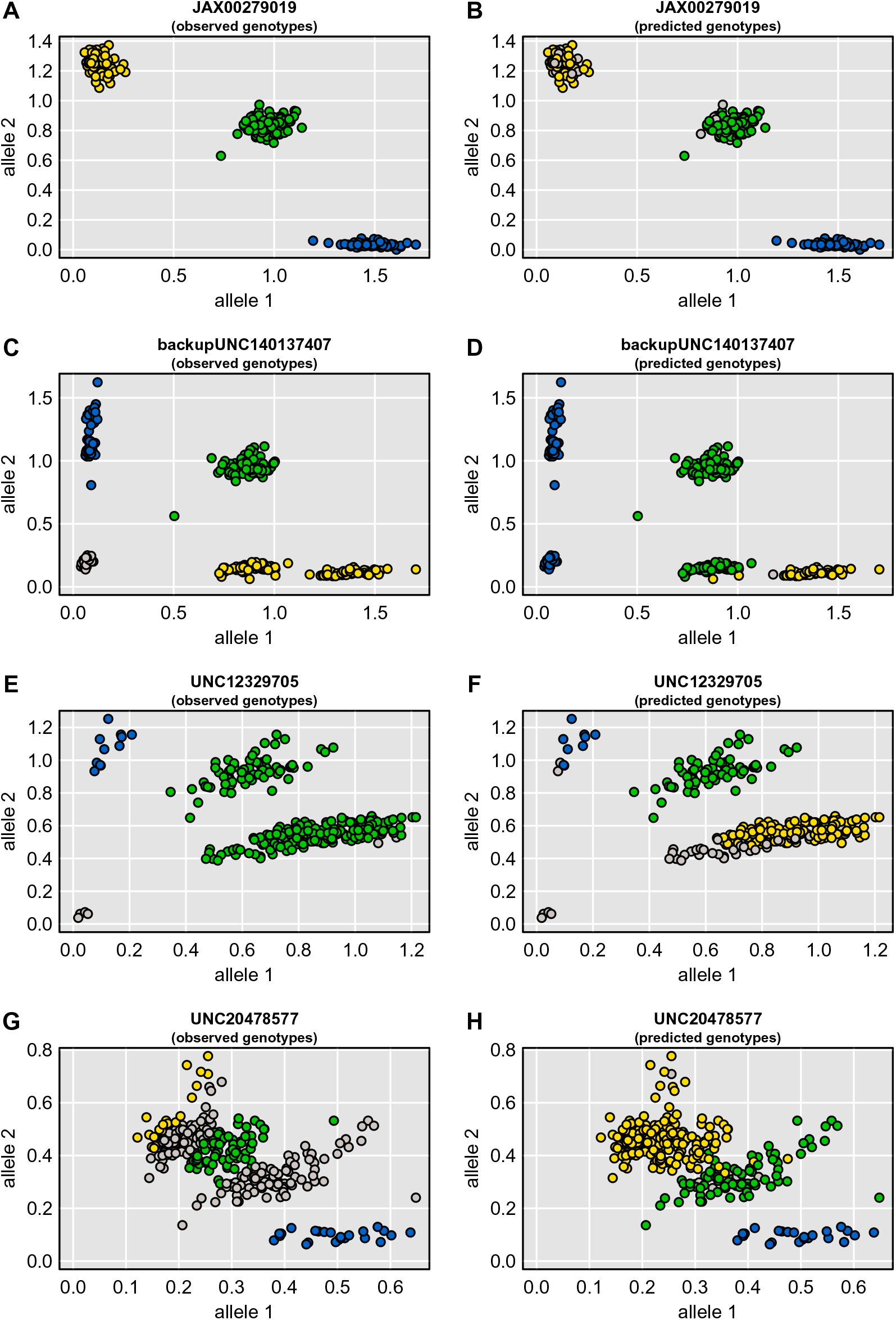
Allele intensity plots for four SNPs. In the left panels, points are colored according to the genotype calls, with yellow and blue being the two homozygotes and green being the heterozygote; gray points were not called. In the right panels, points are colored by the inferred SNP genotypes, given the multipoint marker data and the founders’ genotypes; gray points could not be inferred.

Figures 9C and 9D correspond to a SNP where there is an additional cluster of genotypes, and where the genotype calling algorithm assigned it to the wrong genotype. At this particular SNP, there is a cluster of genotypes that were called homozygotes for the major allele but that are really heterozygotes. This is an example of a variable intensity oligonucleotide (VINO; Didion *et al*. 2012), where there is an additional SNP within the probe that leads to reduced intensity of one allele. For this particular SNP, the founder strains 129Sv/ImJ and PWK/PhJ appear to have null alleles for the array probe (data not shown). Additional examples of this type of SNP are shown in Supplementary Figure S5.

Figures 9E and 9F correspond to a SNP where the genotype calling algorithm made a clear mistake. There are three well-defined clusters of genotypes, but one of the homozygotes got called as a heterozygote: the cluster is green in Figure 9E but yellow (and gray, for missing) in Figure 9F. Figures 9G and 9H correspond to another example of a poorly called SNP, where two of the genotype clusters are not well separated, and the genotype calling algorithm made a mistake in identifying the clusters. Supplementary Figures S6 and S7 show additional examples of mistakes in the genotype calling, either because the genotype clusters are arranged horizontally or are not well separated. Figure S8 contains additional examples of particularly ugly SNPs.

### Effects of data cleaning

High-density SNP data are sufficiently redundant that the presence of a small number of poorly behaved SNPs should have little influence on the results. The hidden Markov model used to reconstruct the DO genomes allows for the presence of genotyping errors and so should smooth over the problem markers. But it is worth checking: if we omit the most poorly behaved markers, how much will the diplotype probabilities change?

Of the 69,339 informative markers, we omitted 325 markers with estimated genotyping error rate > 5%, using a threshold of 2 on the genotyping error LOD score to define a presumed error. We then re-calculated the 36-state diplotype probabilities for all mice.

For each mouse at each marker, we take the sum of the absolute differences between the diplotype probabilities, before and after data cleaning, as a measure of change. This takes values between 0 and 2, with 2 indicating that a complete different set of diplotypes have positive probabilities, after the data cleaning.

There are few changes in the diplotype probabilities. Of the 69,339 markers × 280 individuals, there are just 6,048 sites where the sum of the absolute differences was >1, and just 1,798 where it was >1.5. To illustrate the types of changes that are seen, Supplementary Figure S9 shows the diplotype probabilities, before and after omitting 325 bad markers, for three individuals on chromosome 9. In each case, an apparent recombinant segment gets removed.

## Discussion

Our approach to cleaning genotype data is built around a series of diagnostic visualizations that can help us to identify problematic samples and markers. Identifying problem samples (mix-ups and low quality samples) is arguably the most important outcome of data cleaning. A small number of incorrect samples can impact the power of QTL mapping whereas poorly performing markers generally have little impact on diplotype reconstructions.

The simplest diagnostic for sample quality, the amount of missing data by individual, is also highly effective. In addition to high rates of missing data, low quality samples will display higher rates of genotype calling errors, unexpectedly high numbers of predicted crossover events, and unusual allele frequencies. It is important to verify that samples are not duplicated and that the correct sex is assigned to individuals because these problem will not be captured in other quality control diagnostics.

The proportion of missing data is also a good diagnostic for SNP marker quality. Additional diagnostics include genotyping frequencies, and the estimated proportion of genotyping errors.

In the data we used as an illustration, with 291 DO mice from generations 8 and 11, there were a set of nine samples with ≥ 20% missing genotypes. These also showed excessive crossovers and high genotyping error rates and should be omitted from any analyses. There were also two apparent sample duplicates (one being a male/female pair), and one apparent XO female. Finally, there were four samples with higher-than-normal amounts of missing data and genotyping errors, but these samples looked okay otherwise and probably do not need to be omitted. Decisions about which samples to include or omit should be based on the likely impact on mapping analysis and may depend on factors such as the sample size and the extent and quality of phenotyping data.

Verifying sex and identifying sample duplicates are two steps towards identifying sample mix-ups. If there are phenotypes with strong genetic effects, such as coat color, they may be useful to identify further mix-ups. Particularly useful in this regard are genome-scale phenotype data such as gene expression data, whether by microarrays or RNA-seq, which can perfectly identify individuals (Westra *et al*. 2011). With gene expression data on multiple tissues, identified mix-ups can potentially be corrected (Broman *et al*. 2015).

The vast majority of markers appeared well behaved, but we also found a number of markers with a high proportion of apparent genotyping errors. Omitting these markers had small and relatively isolated effects on the diplotype probabilities. Our approach for identifying problem markers relied on our diplotype reconstructions, and so we may be missing some badly behaved SNPs, and the SNPs we miss may be the ones with the greatest influence on results. But if the overall SNP density is high, and the proportion of badly behaved SNPs is low, our approach should provide reasonable results. While poor quality markers may have little impact on any given study, identifying and annotating these markers may impact other studies that use the same array platform.

We have not discussed the problem of cleaning phenotype data, but this is also important. We would focus on data visualizations, including histograms, plots of traits by time of measurement and/or by mouse identifiers, and scatterplots of traits against one another. These plots may reveal typographical errors in the data or inconsistencies in measurement units. They may also indicate the need for phenotype transformations (such as logarithm or square-root), or they may reveal important batch effects or other covariates that should be taken into account in analyses.

We used R (R Core Team 2018) and R/qtl2 (Broman *et al*. 2019) throughout this work. Another important R package for genotype diagnostics is argyle (Morgan 2015), which provides a variety of diagnostics for SNP genotyping arrays.

We focused on data visualizations to diagnose potential problems, and that is the central tool for data cleaning. Make lots of graphs, focusing on graphs that will reveal anticipated problems but also following up on anything unexpected: Is it a problem with the data, a problem with the sample, or a quirk of biology? Is it ignorable or fixable? What effect might it have on later results? We find interactive data visualizations, such as with R/qtlcharts (Broman 2015), useful in these efforts, particularly for identifying outlier samples in scatterplots.

While we have focused on DO mice, our approach could be applied more generally, to other multiparent populations. The key summary statistics are the proportion of missing genotypes, the average probe intensities for SNPs on the X and Y chromosomes, the proportion of heterozygous SNPs, the estimated number of crossovers, and the estimated genotyping error rate.

## Acknowledgments

This work was supported in part by National Institutes of Health grants R01GM070683 (to K.W.B. and G.A.C.) and R01GM123489 (to Ś.S.).

---

**Figure S1.**
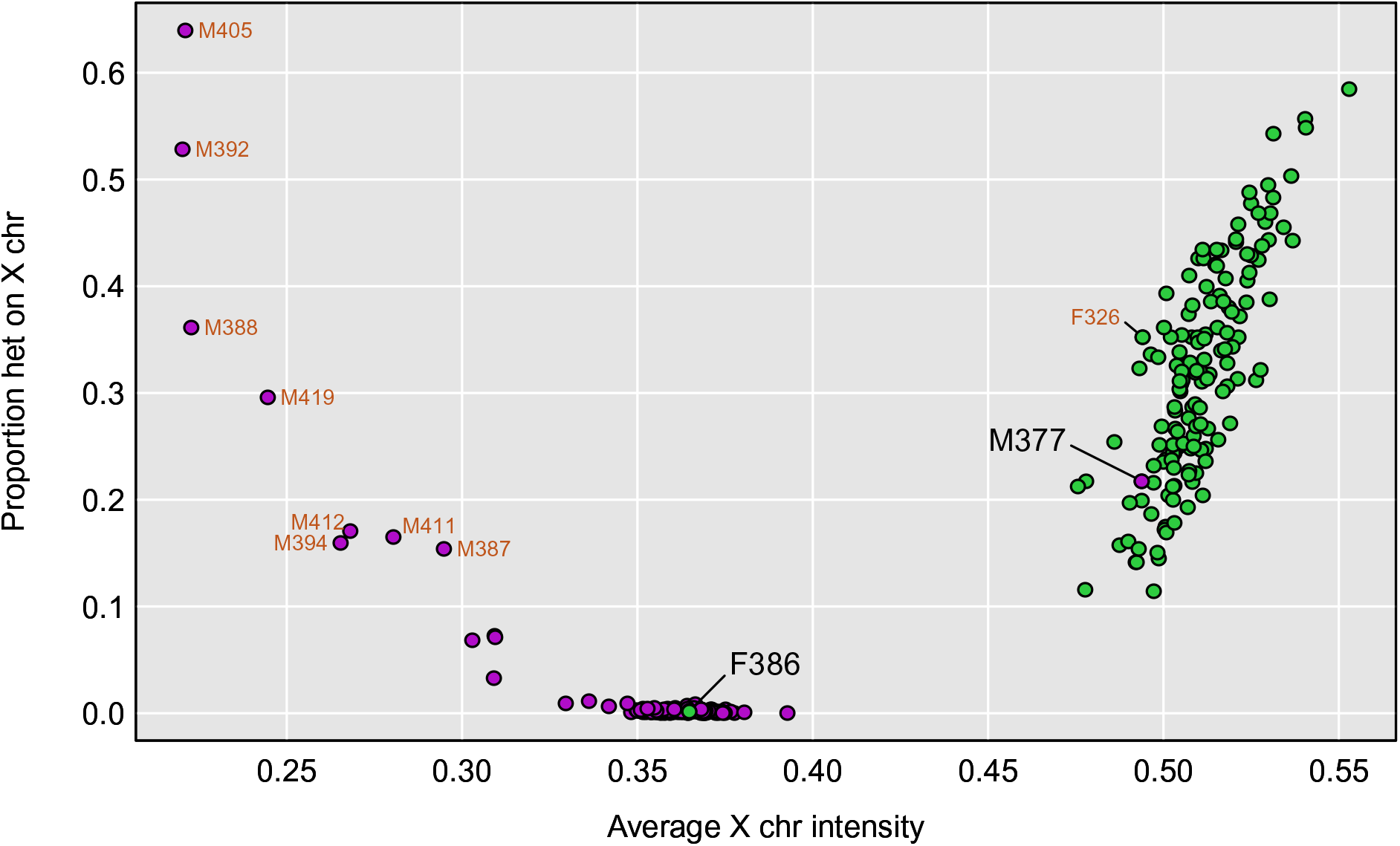
Proportion of heterozygous SNPs on the X chromosome versus the average microarray intensity for markers on the X chromosome, for each mouse. Mice that were nominally male are in purple, while females are in green. Samples with ≥ 20% missing data are labeled in orange. Two other samples of interest are labeled in black: F386 which appears to be XO, and M377 which was nominally male but appears to be XX.

**Figure S2.**
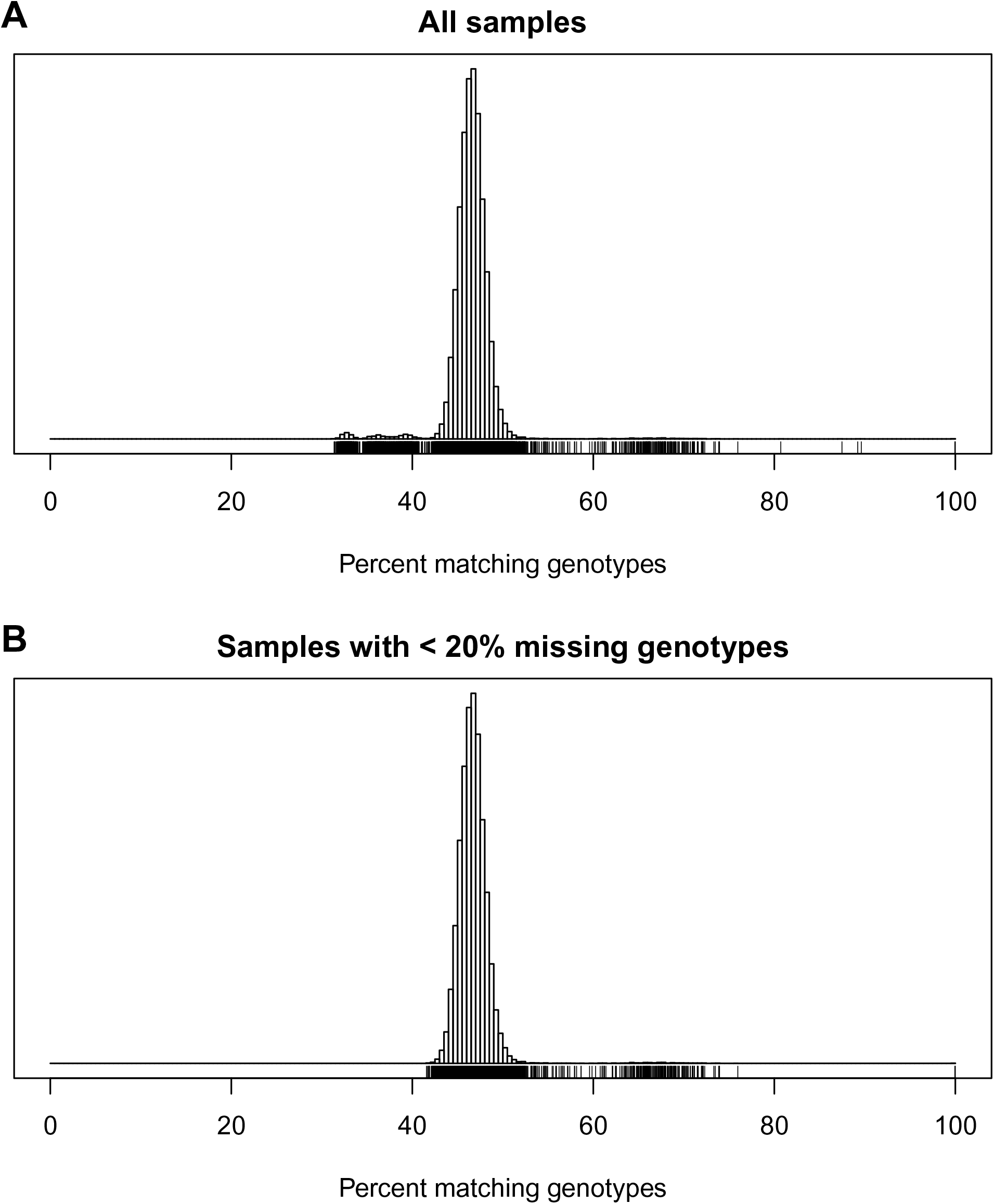
Distribution of percent matching genotypes for pairs of mice. A: all samples. B: samples with < 20% missing genotypes. Tick marks below the histograms indicate the values for individual pairs.

**Figure S3.**
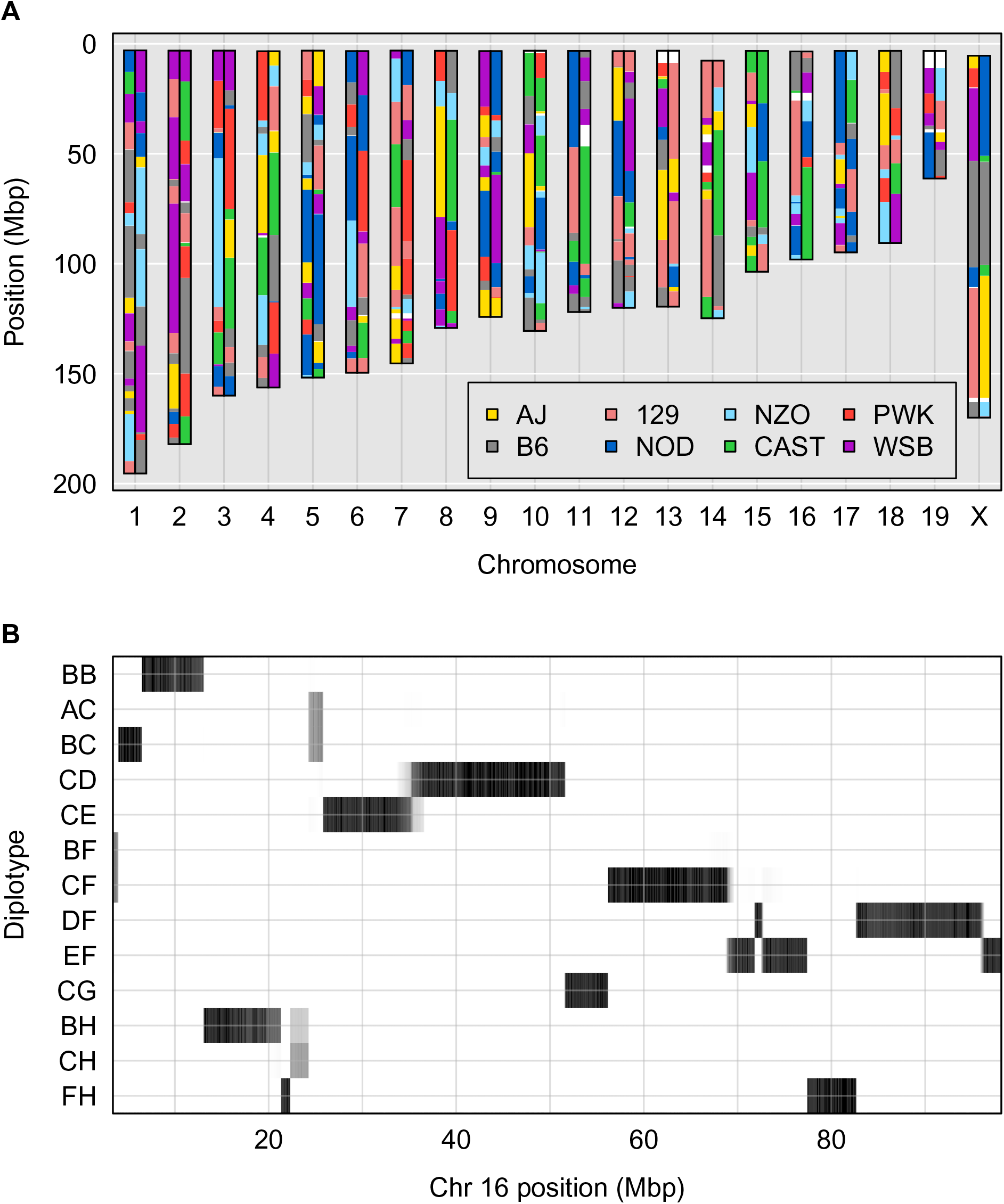
Illustration of genome reconstruction. **A**: Inferred diplotype across the genome for mouse F413, with an arbitrary choice of phase. (For example, if there is a region of homozygosity, the haplotypes above can be swapped relative to the haplotypes below.) White indicates unknown (no diplotype had probability >0.5). **B**: Heat map of the diplotype probabilities for mouse F413 along chromosome 16. Only diplotypes that achieved probability > 0.25 are shown.

**Figure S4.**
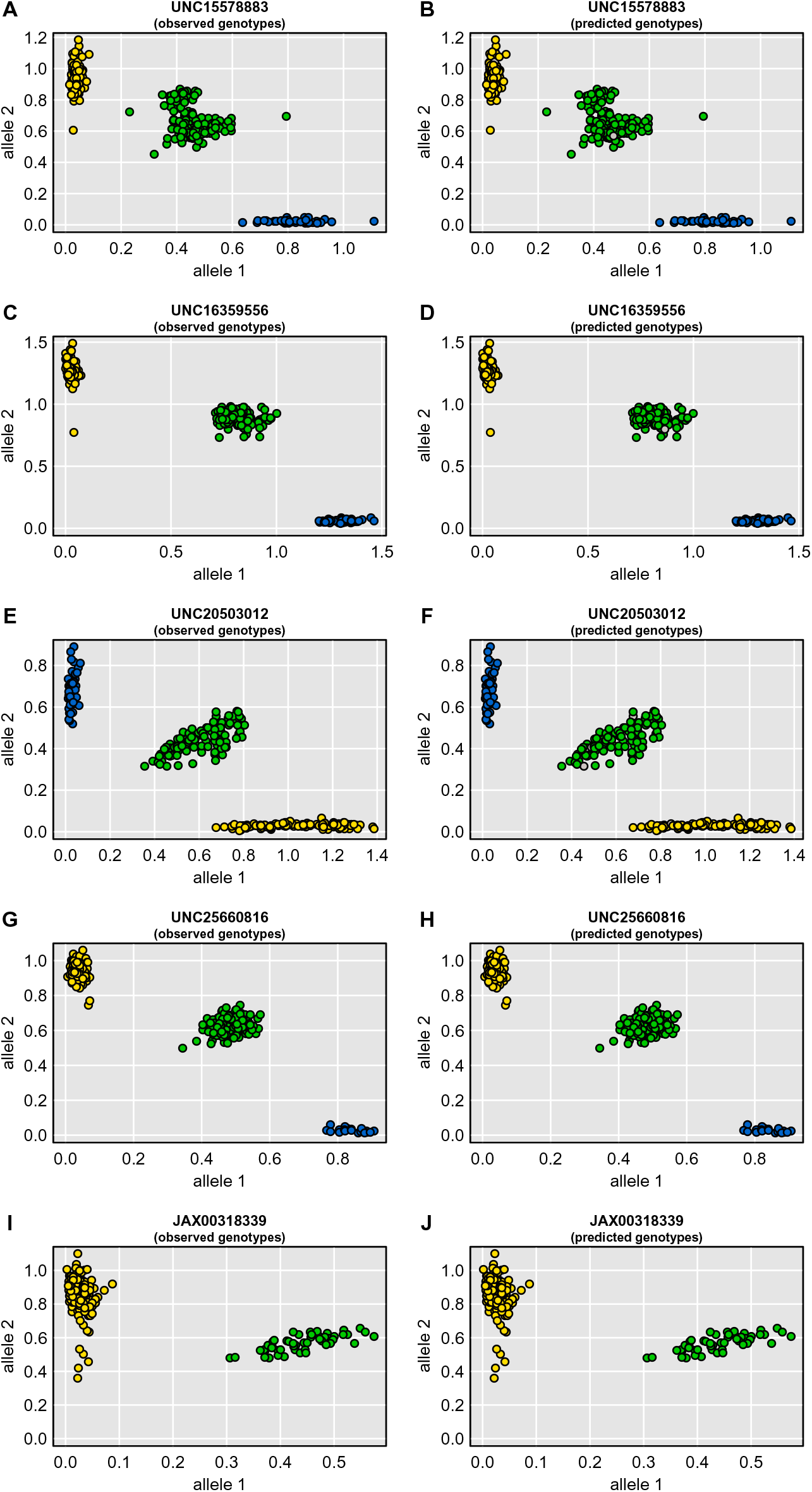
Allele intensity plots for examples of well-behaved SNPs. In the left panels, points are colored according to the genotype calls, with yellow and blue being the two homozygotes and green being the heterozygote; gray points were not called. In the right panels, points are colored by the inferred SNP genotypes, given the multipoint marker data and the founders’ genotypes; gray points could not be inferred.

**Figure S5.**
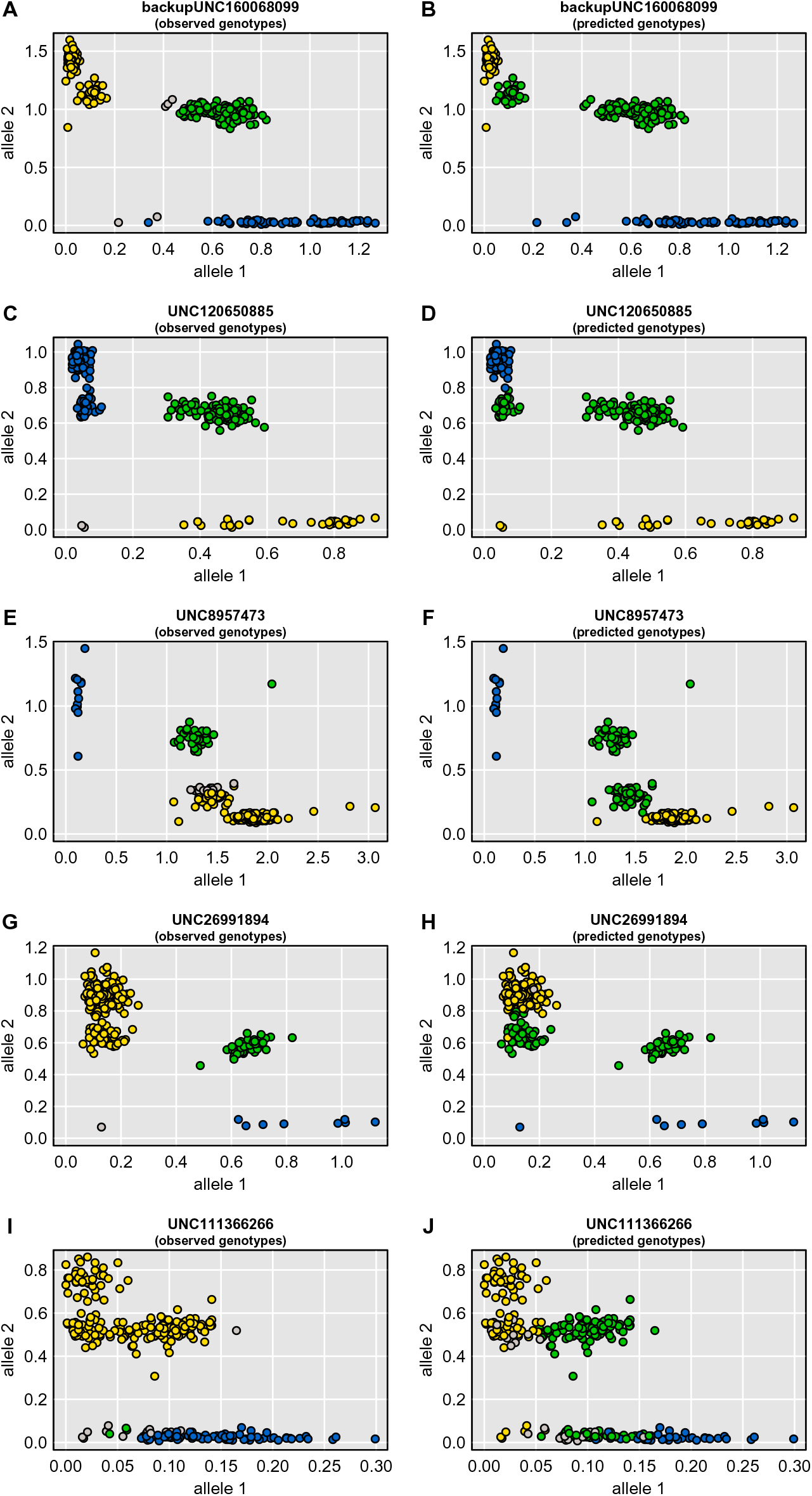
Allele intensity plots for examples of SNPs with an additional genotype cluster. In the left panels, points are colored according to the genotype calls, with yellow and blue being the two homozygotes and green being the heterozygote; gray points were not called. In the right panels, points are colored by the inferred SNP genotypes, given the multipoint marker data and the founders’ genotypes; gray points could not be inferred.

**Figure S6.**
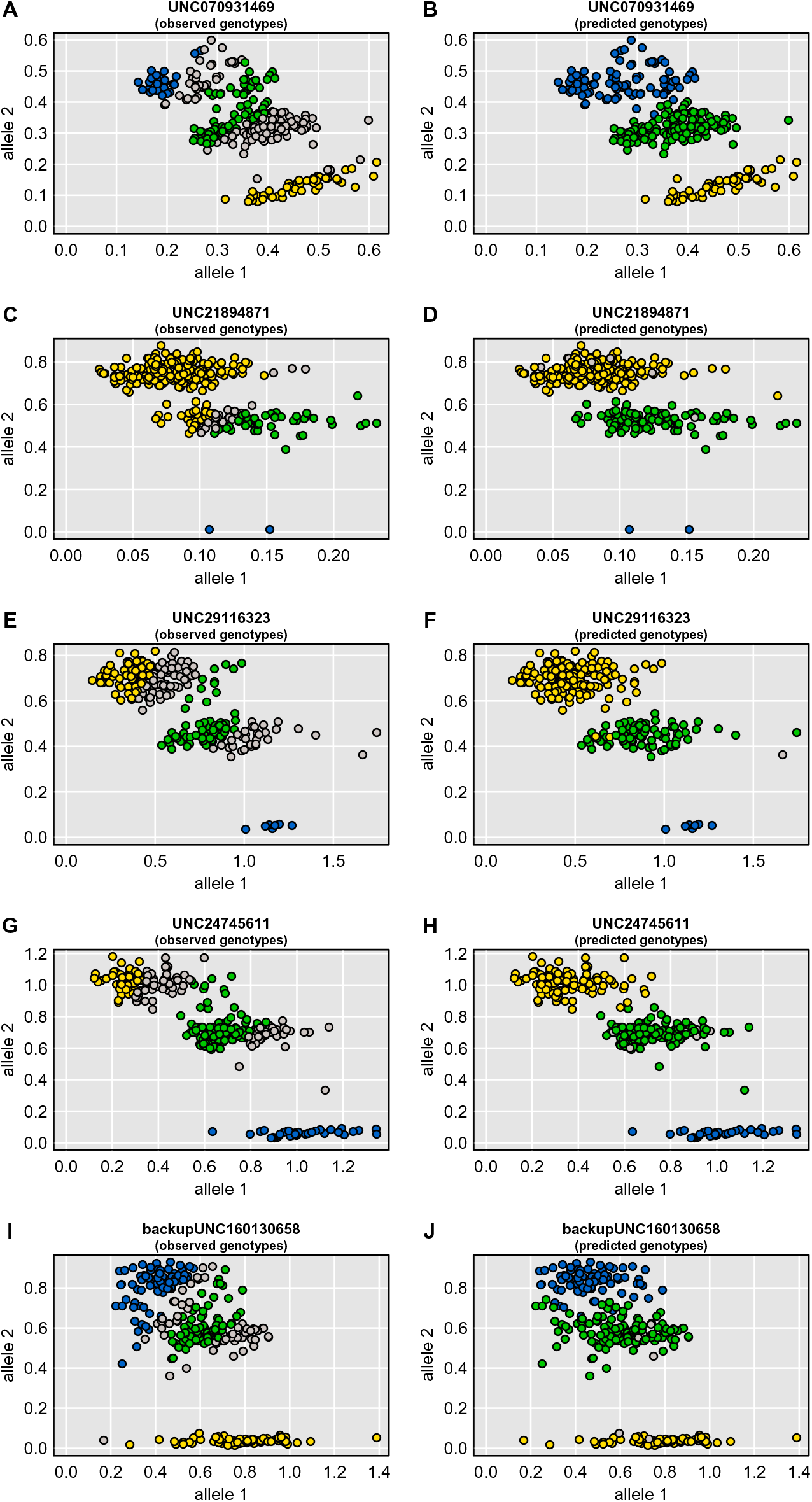
Allele intensity plots for examples of SNPs that are error-prone due to clusters oriented horizontally. In the left panels, points are colored according to the genotype calls, with yellow and blue being the two homozygotes and green being the heterozygote; gray points were not called. In the right panels, points are colored by the inferred SNP genotypes, given the multipoint marker data and the founders’ genotypes; gray points could not be inferred.

**Figure S7.**
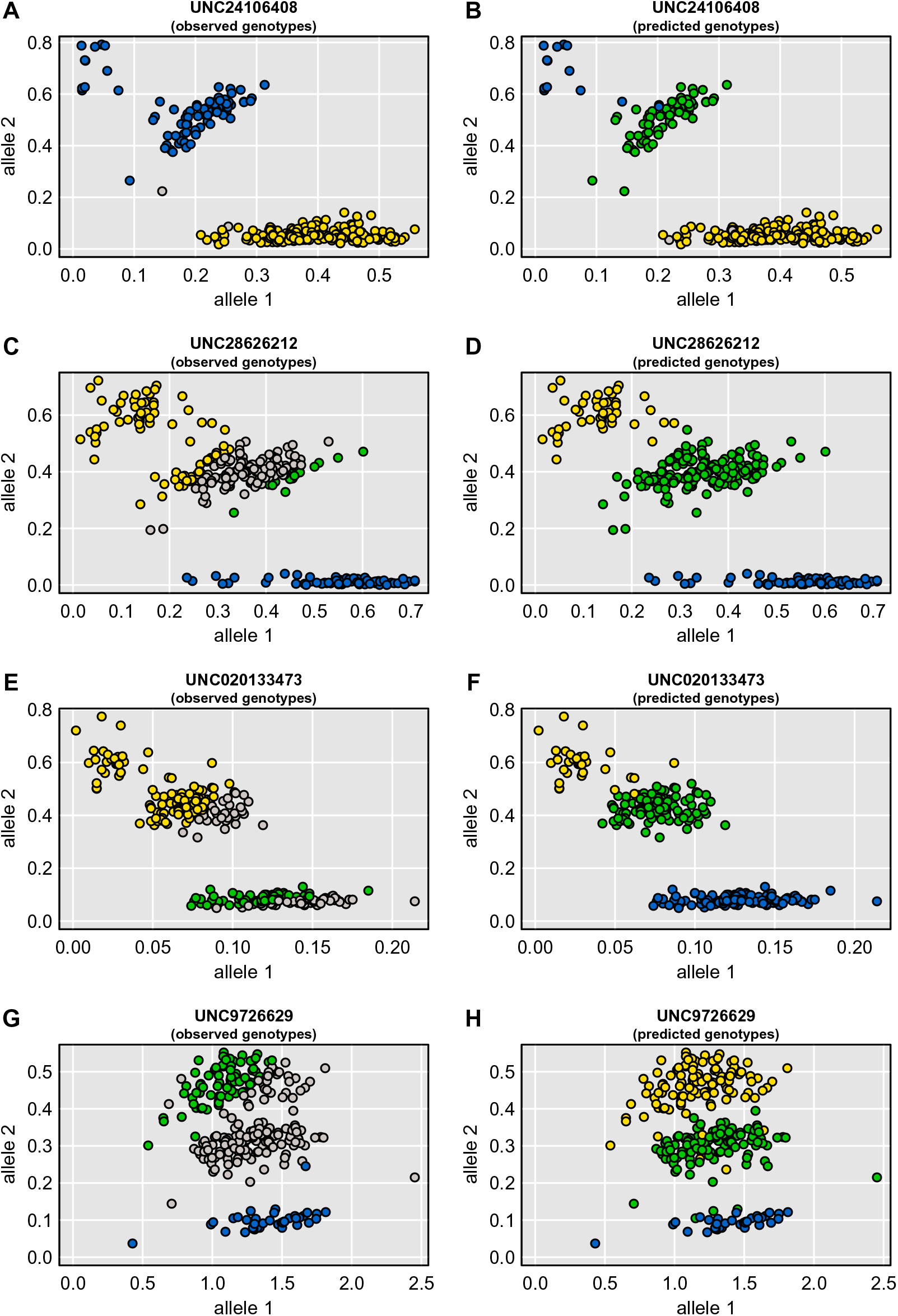
Allele intensity plots for examples of SNPs that have been poorly called. In the left panels, points are colored according to the genotype calls, with yellow and blue being the two homozygotes and green being the heterozygote; gray points were not called. In the right panels, points are colored by the inferred SNP genotypes, given the multipoint marker data and the founders’ genotypes; gray points could not be inferred.

**Figure S8.**
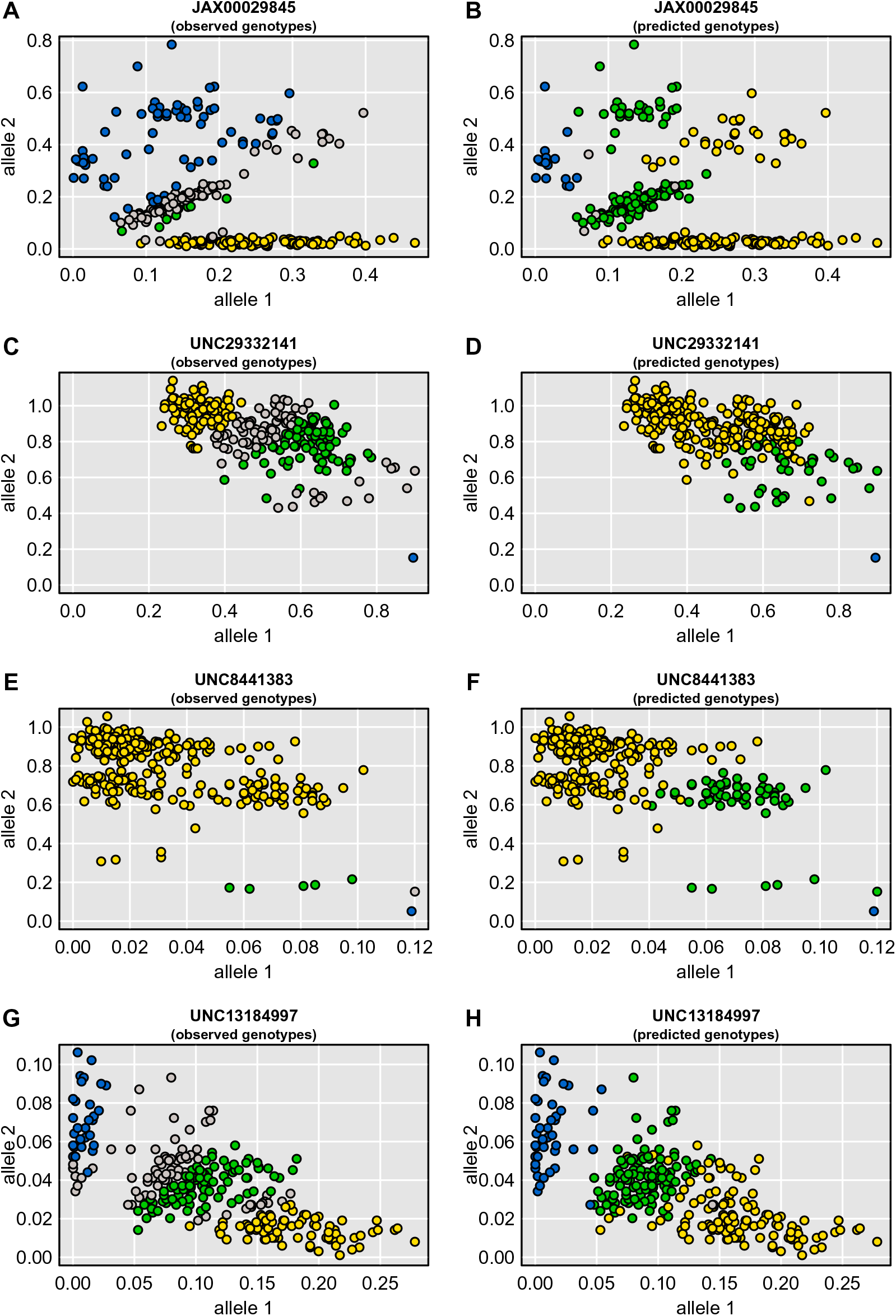
Allele intensity plots for examples of SNPs that are particularly ugly. In the left panels, points are colored according to the genotype calls, with yellow and blue being the two homozygotes and green being the heterozygote; gray points were not called. In the right panels, points are colored by the inferred SNP genotypes, given the multipoint marker data and the founders’ genotypes; gray points could not be inferred.

**Figure S9.**
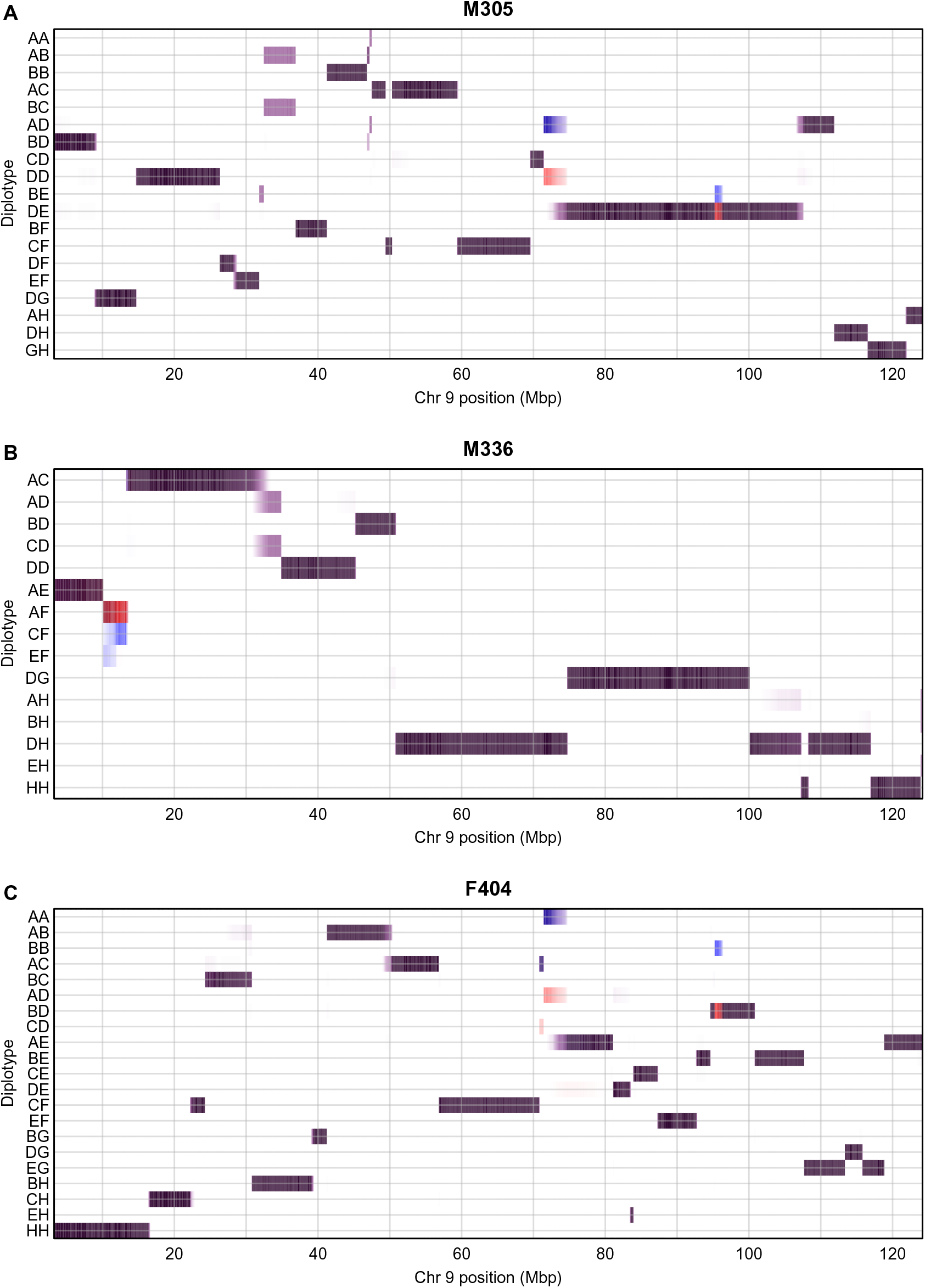
Bivariate heatmaps displaying diplotype probabilities before and after omitting badly behaved markers, for three selected individuals on chromosome 9. Probabilities before and after omitting markers are shown in white/blue and white/red color scales, respectively. Only diplotypes that achieved probability > 0.25 are shown. Dark purple indicates the probability was high both before and after data cleaning, blue indicates high before data cleaning but low after, and red indicates low before but high after.

